# Synthetic G-quadruplex components for predictable, precise two-level control of mammalian recombinant protein expression

**DOI:** 10.1101/2024.09.10.612233

**Authors:** Melinda Pohle, Edward Curry, Suzanne Gibson, Adam Brown

## Abstract

Control of mammalian recombinant protein expression underpins the *in vitro* manufacture and *in vivo* performance of all biopharmaceutical products. However, routine optimization of protein expression levels in these applications is hampered by a paucity of genetic elements that function predictably across varying molecular formats and host cell contexts. Herein, we describe synthetic genetic components that are specifically built to simplify bioindustrial expression cassette design processes. Synthetic G-quadruplex elements with varying sequence feature compositions were systematically designed to exhibit a wide-range of regulatory activities, and inserted into identified optimal positions within a standardized, bioindustry compatible core promoter-5’UTR control unit. The resulting library tuned protein production rates over two orders of magnitude, where DNA and RNA G-quadruplexes could be deployed individually, or in combination to achieve synergistic two-level regulatory control. We demonstrate these components can predictably and precisely tailor protein expression levels in i) varying gene therapy and biomanufacturing cell hosts, and ii) both plasmid DNA and synthetic mRNA contexts. As an exemplar use-case, a vector design platform was created to facilitate rapid optimization of polypeptide expression ratios for difficult-to-express multichain products. Permitting simple, predictable titration of recombinant protein expression, this technology should prove useful for gene therapy and biopharmaceutical manufacturing applications.

## 1. Introduction

Recombinant protein expression in mammalian cells is underpinned by genetic technologies that control flux through the key steps in product biosynthesis, namely gene transcription, mRNA translation and polypeptide translocation (1). In most cases, these processes are still controlled by assemblies of viral and endogenous elements that are generally known to encode reasonably high biosynthetic rates, such as the CMV-IE1 promoter, human alpha-globin 5’ UTR and SV40 3’UTR (2–4). However, increasingly complex protein expression objectives have created a requirement for more sophisticated gene expression control technologies that can precisely tailor transcription and translation rates in molecule- and context-specific manners. These include i) advanced mammalian cell factory engineering strategies, where multiple genes must be expressed at correct stoichiometric ratios to achieve designed phenotypes (5), ii) gene therapy, where target proteins need to be expressed at levels which provide therapeutic efficacy without causing off-target effects (6), and iii) protein biopharmaceutical manufacturing, where the expression levels of multiple polypeptide chains must be optimally balanced to facilitate correct protein folding and assembly (7). Indeed, in the latter example, many promising product molecules are currently considered difficult or impossible to manufacture due to improperly assembled proteins inducing cellular stress responses (8, 9).

Predictable and precise control of recombinant protein expression levels requires design and validation of synthetic genetic component libraries. While synthetic signal peptides can be used to tailor translocation levels, their activity is highly polypeptide-specific, and this lack of orthogonality significantly restricts their routine use in vector design spaces owing to the associated *in vitro* screening burden (10). Conversely, synthetic promoter and UTR elements have been developed to customize transcription and translation levels, but their performance is typically cell-type and cell-context specific, dependent on relative abundances of cognate nucleic-acid binding proteins (11, 12). This necessitates design of cell host specific genetic component libraries, which are only currently available for a handful of mammalian cells (13–15). Accordingly, for most mammalian protein expression applications, including the vast majority of cell type specific gene therapies, it is currently intractable to design genetic construct assemblies that encode optimal biosynthetic rates, owing to a paucity of appropriate validated regulatory components.

Secondary structure-based control elements represent a promising, and relatively unexplored, means to achieve sophisticated mammalian protein expression control. Nucleic acid strands can fold into a variety of stable structures, including loops, hairpins, helices, and pseudoknots. When present in appropriate positions within promoters and UTRs their formation can sterically hinder the binding and progression of transcriptional and translational apparatus, thereby facilitating biosynthetic rate tuning by selecting elements with varying inhibitory activities (e.g. varying thermodynamic stabilities) (16). An elegant example of designing secondary structure components to tailor mammalian protein expression for bioindustrial applications is the study by Eisenhut *et al.*, where 25 distinct RNA hairpins were created by varying GC content, minimum free energy and positioning relative to the 5’ cap (17). This library was used to vary translation rates over two orders of magnitude in common mammalian bioproduction cell hosts, exemplifying the utility of secondary structure elements to solve complex protein expression challenges. However, the available regulatory component toolbox remains relatively sparsely populated, and is particularly lacking technologies that can i) predictably tune transcription levels and, accordingly, ii) enact two-level protein expression control by facilitating simultaneous titration of both transcription and translation rates.

G-quadruplexes (G4s) are four-stranded secondary structures that form from guanine-rich nucleic acid sequences, comprising multiple G-tetrads (a square planar arrangement of four guanine bases interconnected through Hoogsteen hydrogen bonding) (18). Prevalent in both the human genome and transcriptome, they form in DNA and RNA molecules to regulate transcription and translation rates (19–21). While some G4s have been shown to upregulate protein expression, e.g. via transcription factor recruitment (22, 23), when appropriately positioned within promoters and 5’ UTRs they can act to reduce protein biosynthesis by disrupting the function of general transcriptional or translational machinery (24–26). Although hundreds of discrete G4s have been experimentally validated, mainly those that regulate expression of oncogenes such as c-MYC and NRAS (27–29), these endogenous components cannot be easily applied to bioindustrial applications as their functionality has only been profiled within the context of natural human promoters and UTRs (e.g. 30–33) which typically display unpredictable or undesirable expression dynamics (16, 34, 35). Moreover, they are usually studied individually, where information on their relative activities is not generally available, preventing their use as amalgamated libraries where elements can be rationally selected and combined to achieve user-defined protein expression rates. Accordingly, harnessing the potential of G4s for cell-agnostic two-level precision protein expression control requires bottom-up creation of built-for-purpose synthetic libraries, comprising well-characterized elements that are designed and validated to function within standardized industrially relevant promoter-UTR contexts.

In this study we develop the first synthetic G4 element library designed specifically for use in high-value bioindustrial applications. Comprising complementary DNA and RNA G4 components operating at optimized positions within a standardized genetic architecture, this technology enables precise two-level control of mammalian protein expression over two orders of magnitude. Validated to perform predictably in varying cell types and molecular formats (i.e. plasmid DNA, synthetic mRNA, multi-chain products) these synthetic G4 controllers could be utilized to predictably fine-tune recombinant protein expression levels in *in vitro* biomanufacturing and *in vivo* biomedical applications.

## 2. Materials and Methods

### Expression cassette and vector construction

The bioindustrial protein expression control unit (BPCU) was constructed by fusing a minimal 41 bp hCMV-IE1 core promoter (GAN M60321.1, nucleotides 1110 – 1150) to a 57 bp proprietary AstraZeneca 5’ UTR (36). BPCU was inserted upstream of the gene of interest (containing an SV40 polyA 3’ UTR (GAN LT727517.1, nucleotides 1449–1676)) and downstream of the hCMV-IE1 proximal promoter (GAN M60321.1, nucleotides 517–1109) in reporter vectors encoding SEAP, eGFP, mAbT HC, and mAbT LC expression (reporter vector *de novo* synthesized by Geneart (Regensburg, Germany)). G4 elements (listed in Supplementary Table S1) were inserted into BPCU by site-directed mutagenesis using the Q5® Site-Directed Mutagenesis Kit (New England Biolabs, USA). Plasmid templates for *in vitro transcription* reactions were generated by cloning appropriate RNA G4-eGFP sequences into pRNA128A (37) using NheI and NotI (New England Biolabs, USA) restriction sites. Vectors for stable transfections were created by inserting appropriate DNA G4-eGFP sequences into pMCS-Gaussia Luc (Thermo Fisher Scientific, USA) using KpnI and NotI restriction sites. All constructed plasmids used in this study were validated via DNA sequencing and purified to transfection-grade quality using the QIAGEN Plasmid Plus Midi kit (QIAGEN, USA).

### Cell culture

CHO-K1 derived host cells (AstraZeneca, UK), hereby referred to as CHO cells, and CHO-S (Thermo Fisher Scientific, USA) cells were routinely cultivated in CD-CHO medium (Thermo Fisher Scientific, USA) supplemented with 6 mM L-glutamine and 8 mM L-glutamine, respectively. Expi293F cells (Thermo Fisher Scientific, USA), hereby referred to as HEK cells, were routinely cultivated in Freestyle™ 293 expression medium (Thermo Fisher Scientific, USA). Cultures were maintained at 37 °C, 5% CO_2_ with 140 rpm orbital shaking. CHO cells were passaged every 3-4 days at a seeding density of 0.2 x 10^6^ cells/ml, CHO-S cells were passaged every 2-3 days at a seeding density of 0.2 x 10^6^ cells/ml and HEK cells were passaged every 3-4 days at a seeding density of 0.3-0.4 x 10^6^ cells/ml. HepG2 cells (ATCC HB-8065) were cultured in RPMI 1640 medium (Thermo Fisher Scientific, USA) supplemented with 10% fetal bovine serum (FBS) and maintained at 37 °C under 5% CO_2_. Cell concentration and viability were measured with the VI-CELL BLU automated cell counter and cell viability analyzer (Beckman Coulter, USA).

### Transient transfection

CHO cells were transiently transfected by electroporation using the Amaxa Nucleofector system in combination with the SG Cell Line 96-well Nucleofector® Kit (Lonza, Switzerland). 1.86 x 10^6^ cells per well were transfected with 750 ng plasmid DNA or 400 ng mRNA following manufacturer’s instructions. Transfected cells were transferred to a 24 shallow-well plate and incubated in a total volume of 750 μl for 48 h at 37 °C, 5% CO_2_ at 240 rpm orbital shaking. IgG1 (mAbT) LC- and HC-encoding plasmids were transfected at ratios maintaining equal HC:LC gene copy numbers, and incubated for five days with a 10% (v/v) feed added 48 h post-transfection containing 1:1 (v/v) Efficient Feed A (Thermo Fisher Scientific, USA) and Efficient Feed B (Thermo Fisher Scientific, USA). HEK cells were transiently transfected with PEIpro® (Polyplus, France), whereby 1.5 x 10^6^ cells were transfected with 600 ng plasmid DNA mixed with 1.8 μl PEIpro® (1:3 ratio of DNA to PEI) following manufacturer’s instructions. Cells were incubated in a 24 shallow-well plate in a total volume of 750 μl for 48 h at 37 °C, 5% CO_2_ at 240 rpm orbital shaking. HepG2 cells were transfected with Lipofectamine™ 3000 Transfection Reagent (Thermo Fisher Scientific, USA) according to manufacturer’s instructions in a 24 well plate scale and incubated for 48 h.

### Stable transfection

CHO-S cells were stably transfected by electroporation using the Amaxa Nucleofector system in combination with the Cell Line Nucleofector® Kit V (Lonza, Switzerland). 5.0 x 10^6^ cells per cuvette were transfected with 2 µg plasmid and incubated in T75 flasks at 37 °C under 5% CO_2_ in 13 ml CD-CHO medium supplemented with 8 mM L-glutamine. 48 h post-transfection the medium was changed to CD-CHO selection medium including 8 mM L-glutamine, 10 µl/ml Puromycin and Anti-Clumping agent (Thermo Fisher Scientific, USA) at a 1:250 (v/v) dilution. Once cultures reached > 80%, they were transferred from static to shaking conditions and passaged every 3 days until they reached a viability of > 95% at which point they were cryopreserved.

### Recombinant protein quantification

eGFP protein expression was quantified 48 h post-transfection using the SpectraMax iD5 Microplate Reader (Molecular Devices, USA) with 485 nm excitation and 535 nm emission. Transfection efficiency was determined using the Countess II FL Automated Cell Counter (Thermo Fisher Scientific, USA). SEAP expression was quantified 48 h post-transfection using the Secreted Alkaline Phosphatase Reporter Gene Assay Kit (Cayman Chemical, USA) according to manufacturer’s instructions. For IgG1 mAb protein titer quantification, the ELISA quantification kit for human immunoglobulins G (RD-Biotech, France) was used in accordance with the manufacturer’s protocol five days post-transfection. All assays were performed using the SpectraMax iD5 Microplate Reader.

### Circular Dichroism

DNA oligos (Integrated DNA Technologies, USA) were prepared at 5 µM in 50 mM Tris-HCl (pH 7.5) with 100 mM KCl. The samples were denatured at 95 °C for 5 min and then cooled down 0.2 °C/min to 20°C. Circular dichroism (CD) experiments were performed at 25°C using a JASCO J-810 spectropolarimeter (JASCO, USA) in 1 mm quartz cuvettes. CD scans were taken from 320 nm to 200 nm at 100 nm/min.

### *In vitro* transcription

Plasmid templates for *in vitro* transcription (IVT) were linearized with BsaXI and purified by ethanol precipitation. IVT reactions used the HiScribe® T7 High Yield RNA Synthesis Kit (New England Biolabs, USA) following the manufacturer’s instructions and 3′-O-Me-m^7^G(5′)ppp(5′)G RNA cap (New England Biolabs, USA) was used as the cap structure analog. Samples were digested with DNase I (Thermo Fisher Scientific, USA) for 15 min at 37°C and immediately purified using the Monarch® RNA Cleanup Kit (New England Biolabs, USA). RNA samples were diluted to the desired working concentration and stored at -80 °C prior to use.

### Chemical ligands

CHO-S cell pools were seeded at 1 x 10^6^ cells/ml in 24 shallow-well plates prior to ligand addition. Phen-DC3 (Selleck Chemicals, USA), Pyridostatin (Merck, Germany), TMPyP4 (Selleck Chemicals, USA) and 360A (ApexBio Technology, USA) were added to a final concentration of 25, 5, 25, and 20 µM, respectively (concentrations optimized to maintain cell viabilities > 90%), and cells were maintained for 48 h at 37 °C, 5% CO_2_ with 140 rpm orbital shaking. Phen-DC3 and 360A were dissolved in DMSO and Pyridostatin and TMPyP4 in water.

### Statistical analysis

A one-way ANOVA was performed to evaluate statistical differences with statistical significance being defined as p ≤ 0.05 (* = p ≤ 0.05, ** = p ≤ 0.01, *** = p ≤ 0.001, **** = p ≤ 0.0001). Tukey’s post-hoc test was used to determine statistical differences to other groups and the Dunnett’s post-hoc test was used to determine statistical differences to a specific control. Statistical tests were performed using GraphPad Prism 10 software.

## 3. Results and Discussion

### 3.1 Optimization of DNA and RNA G4 element positioning within a standardized genetic chassis

The human CMV-IE1 core promoter is the most widely utilized regulatory element for controlling mammalian recombinant gene transcription in high-value applications, including gene therapy and biopharmaceutical production, and has been shown to function in conjunction with varying proximal promoter partners (38–40). While a range of human and mouse sequences are available for use as translational controllers, to facilitate deployment in biomedical settings, we combined the CMV-IE1 core with a minimized 5’ UTR that has been specifically developed and validated for use in bioproduction processes (AZ 5’ UTR). This 98 bp bioindustrial protein expression control unit (BPCU) is an ideal genetic chassis to house regulatory G4s that are designed to control transcription and translation rates by interfering with the activity of general machinery components (Fig 1A). Given that G4 function is position-dependent (29, 41), we first sought to identify the optimal locations to place DNA and RNA G4s within this sequence context. As shown in Figure 1B, model DNA (c-MYC) and RNA (NRAS) G4s were each individually inserted into 7 and 6 distinct sites respectively within the BPCU in eGFP-reporter plasmids harboring a CMV-IE1 proximal promoter, selected to drive a high protein expression set-point that can be tuned down by G4 activity (40, 42, 43). Positions were chosen whereby G4 formation should theoretically interfere with binding and movement of biosynthetic machinery, including near key regulatory binding sites such as the TATA box, INR and Kozak sequence (44, 45). DNA and RNA G4-encoding sequences were inserted into the template and coding strand respectively to facilitate formation of secondary structures at the DNA and RNA level (i.e. RNA G4s form in the 5’ UTR of eGFP-mRNA molecules, Fig 1A).

**Figure 1:**
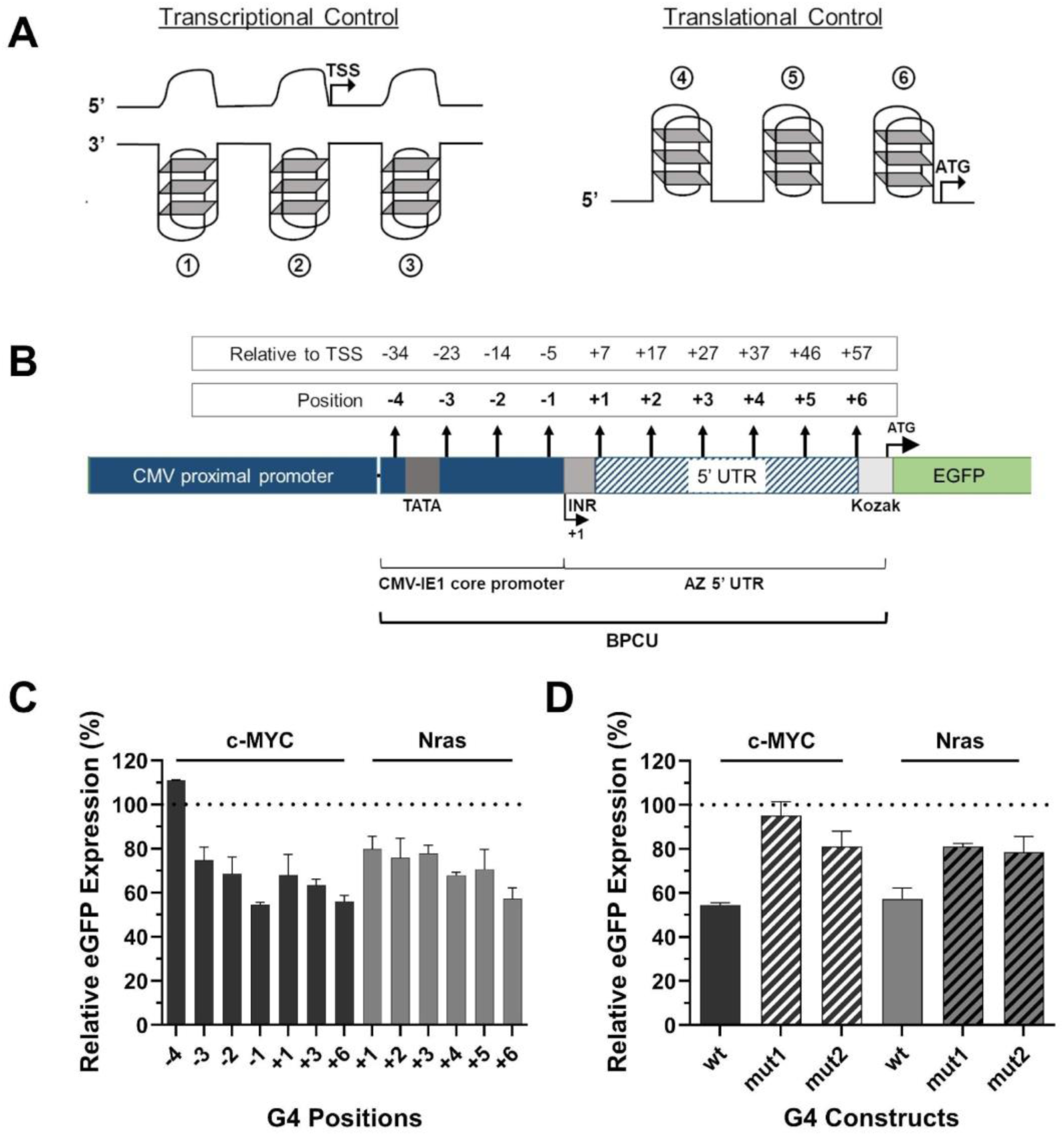
**A)** G-quadruplexes (G4s) in core promoter and 5’ UTR elements can regulate protein expression rates by disrupting the binding (1, 4), movement (3, 5) and activity (2, 6) of transcription and translation initiation complexes. **B)** G4 components were inserted into various positions in a bioindustrial protein expression control unit (BPCU) in eGFP-reporter plasmids harboring a CMV-IE1 proximal promoter. CHO cells were transfected with **C)** reporter vectors containing c-MYC (DNA) and Nras (RNA) G4 elements at different sites within the BPCU, and **D)** eGFP-reporters containing wild-type or mutated versions (see Supplementary Table 2) of c-MYC and Nras G4s in positions -1 and +6 respectively. eGFP expression was quantified 48 h post-transfection. Data are expressed as a percentage of the production exhibited by the control CMV-IE1 proximal promoter-BPCU (ΔG4) construct (dotted line). Values represent the mean + *SD* of three independent experiments (*n* = 3, each performed in triplicate). TSS – transcriptional start site.

G4-containing reporter plasmids were independently transfected into Chinese Hamster Ovary (CHO) cells, selected as a model mammalian cell line due to it being the dominant cell factory for biopharmaceutical production, and eGFP expression levels were quantified 48 h later. These data show that G4s are functional within the context of the BPCU, where protein expression was substantially downregulated by G4s acting at both the DNA and RNA level, as compared to the unengineered control construct (i.e. BPCU without G4 insertion, Fig 1C). Indeed, G4 insertion had a negative regulatory effect on eGFP expression in 12 of 13 tested sequence positions, only being non-functional in the DNA position that was furthest from the transcriptional start site. G4s exerted greatest inhibitory effect when located directly upstream of the transcriptional and translational start sites, in both cases reducing eGFP expression by ∼45% compared to the standard BPCU. These findings are consistent with computational analyses showing that endogenous G4s are preferentially located near transcription start sites (46). Previous studies have found that positional effects on RNA G4 function are context-specific, depending on the 5’ UTR sequence (29, 41). Ultimately, position-optimization studies should be conducted for any new genetic chassis given that G4 inhibitory activity may be impacted by both the relative location of other regulatory elements, and the nucleotide-composition of flanking sequences either side of the insertion site that can affect G4 formation kinetics (47, 48). Within the context of the BPCU, we concluded that positions -1 and +6 were optimal locations for DNA and RNA elements respectively, where inhibitory effects of ∼45% facilitates subsequent design of varying strength G4 components that can titrate protein expression levels both above and below this set-point.

To validate that decreases in eGFP expression were caused by secondary structure-based biosynthesis inhibition, we inserted mutated versions of the c-MYC and NRAS G4 elements into positions -1 and +6 in the BPCU. Mutants were designed to maintain high sequence similarity and equal length to originator elements, while substantially decreasing their propensity to form G4 structures (predicted using QGSR mapper (49) – see Supplementary Table 2). As shown in Figure 3D mutated motifs reduced eGFP expression by 5-20%, compared to the standard BPCU. A generic insertional effect is unsurprising, as previous studies have shown that i) mutated G4 motifs have a similar impact in other genetic contexts (50), and ii) small changes to mammalian core promoters and 5’ UTRs, including the CMV-IE1 core, typically reduce protein expression levels (51). However, given that wild-type G4 sequences had a substantially greater inhibitory effect (> 2.5x) than disabled versions containing a minimal number of G-to-A mutations, we inferred that these elements were functioning via secondary structure-based interference of transcription and translation processes. Accordingly, we concluded that this genetic architecture (i.e. the BPCU with optimized DNA and RNA G4 insertion points) could be employed to create a synthetic G4 component library for two-level precision control of recombinant protein expression.

### 3.2 Synthetic G4 components facilitate predictable, tunable control of recombinant protein expression

G4s have a simple core structure, consisting of alternating runs of guanine tracts (G-tracts) and separating loops, with four possible key input design parameters, namely loop length, G-tract length, loop nucleotide composition and G-tract number (Fig 2A). Using a small number of exemplar elements, it has previously been shown that functional synthetic RNA G4s with different regulatory properties can be created by varying one or more of these design features (41). Accordingly, we rationalized that a component library capable of precisely tailoring protein expression levels from 0 to 100% relative to the standard BPCU could be derived by designing DNA and RNA G4s with varying input parameter combinations. Such G4 variants should exhibit variable propensities to both form and maintain secondary structures within the standardized BPCU genetic chassis, facilitating development of regulatory elements with different inhibitory activities (i.e. varying abilities for steric hindrance-mediated downregulation of transcription and translation rates). Based on previous studies showing loop nucleotide composition to have a relatively minimal impact on G4 thermal stability (52), we adopted two distinct design strategies, namely i) keeping G-tract number constant (N = 4) and varying G-tract length (N = 2-6) and loop length (N = 1-5), and ii) keeping loop length constant (n = 2) and varying G-tract length (N = 2-6) and number (N = 4-7). In both cases we applied a full-factorial design and tested each element at the DNA and RNA level, resulting in a library of 80 distinct G4 components, containing loop sequences that maintained equal A:T:C nucleotide ratios within each construct (all synthetic G4 sequences are provided in Supplementary Table 1).

**Figure 2:**
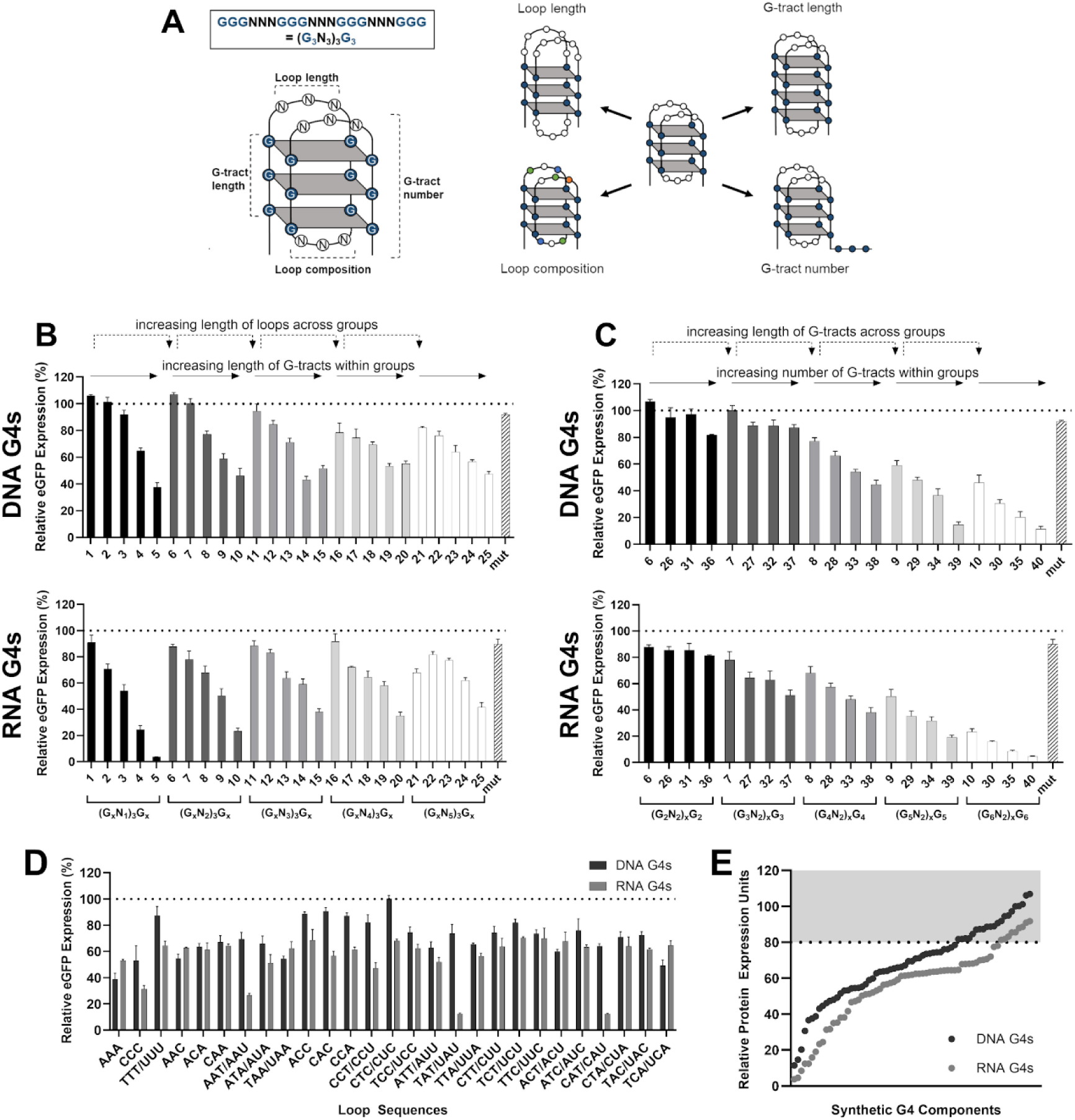
**A)** Core G-quadruplex (G4) structure, highlighting four key input design parameters. **B-C)** Synthetic G4 components (listed in Supplementary Table 1) with varying sequence feature compositions were inserted into positions -1 (DNA) and +6 (RNA) respectively in the BPCU (see Fig. 1) downstream of a CMV-IE1 proximal promoter in eGFP-reporter vectors. **D)** The loop sequence within G4 motif 13 ((G_4_N_3_)_3_G_4_) was varied at both the DNA and RNA level. In B-D, CHO cells were transfected with eGFP-reporter plasmids prior to protein quantification after 48 h. Data are expressed as a percentage of the production exhibited by the control CMV-IE1 proximal promoter-BPCU (ΔG4) construct (dotted line). Values represent the mean + SD of three independent experiments (n = 3, each performed in triplicate). **E)** The library of synthetic G4 components encodes a wide range of recombinant protein expression set points; the gray shaded area represents the generic insertional effects observed when mutated G4s are inserted into the BPCU.

Synthetic G4s were individually inserted in the identified optimal positions in the BPCU (DNA elements in position -1, RNA elements in position +6, Fig 1) downstream of a CMV-IE1 proximal promoter in eGFP-reporter vectors and transfected into CHO cells. Quantification of eGFP expression levels 48 h post-transfection showed that engineered G4s displayed a wide range of inhibitory activities, driving expression levels between 11 – 107% and 4 – 90% at the DNA and RNA level respectively, compared to the standard BPCU control (Fig 2B-C). We note that four new mutant variants of varying sizes exhibited the same generic insertional effect as seen previously, reducing eGFP expression by 5 – 20% (data not shown). Accordingly, we concluded that only those components that reduced expression by > 20%, 57 synthetic G4s in total, were exerting secondary structure-based control of protein biosynthesis. The inhibitory activity of G4 sequences was highly correlated at the DNA and RNA level (Pearson’s correlation coefficient, r = 0.85, p = < 0.0001) identifying that discrete designs behave similarly when utilized as transcriptional or translational controllers. As depicted in Figure 2B-C, G4 performance could be predictably tailored by varying any of the selected input parameters, where inhibitory activities were particularly correlated with increasing G-tract length and number. Indeed, multiple linear regression modelling confirmed that G4 inhibitory activity could be accurately explained as a function of these three features (DNA elements model: R^2^ = 0.85 p = < 0.0001; RNA elements model: R^2^ = 0.87 p = < 0.0001). These findings correlate with previous oligonucleotide-based studies that have shown G-tract length, G-tract number and loop length to be critical determinants of G4 thermal stabilities (41, 52–54). To our knowledge, this represents the first model linking synthetic G4 sequence feature composition to protein expression regulatory function. We reasoned that the ability to accurately explain the performance of these synthetic components as a function of their sequence parameters will enhance both the confidence with which they can be applied to new applications (i.e. as unknown features are unlikely to cause unpredictable changes to activities) and the ability to (re)engineer future elements with defined characteristics.

We predicted that loop nucleotide content would have a negligible impact on G4 performance (52) and accordingly that these ‘non-functional sections’ may provide opportunities to achieve other design objectives such as enhanced sequence efficiency (e.g. by incorporating other regulatory motifs), tailored immunostimulatory effects (55, 56) and optimized GC-content (57). To evaluate this, we varied loop sequence within a standard G4-architecture (motif 13 - (G_4_N_3_)_3_G_4_), where all 27 possible 3-nucleotide permutations (using A, T and C nucleotides only) were individually tested (e.g. ATC: GGGATCGGGATCGGGATCGGG). Quantitative evaluation in CHO cells confirmed that loop composition generally has minimal impact on component inhibitory activity, where 17 of 27 DNA and 23 of 27 RNA elements drove eGFP expression levels within one standard deviation of the mean (Fig 2D). Notably, the three motifs that had the greatest impact on eGFP levels, RNA loops of AAU, UAU and CAU, all introduced alternative start codons, highlighting the requirement for multi-parameter/objective optimization approaches when designing new synthetic nucleotide parts (58). Discounting these three sequences (and any components that reduced eGFP expression by < 20% relative to the standard BPCU) based on the assumption that they encode undesirable variation in translational start sites, our final library comprises 101 distinct regulatory elements (by far the largest library of synthetic G4 controllers described to-date) that facilitate predictable two-level precision control of recombinant protein expression over two orders of magnitude within a standardized industrially relevant genetic chassis (Fig 2E).

### 3.3 Mechanistic dissection of synthetic G4 component functionality and performance

To facilitate their adoption in bioindustrial applications, we next sought to obtain detailed mechanistic information on the function and performance of our synthetic G4 components. Firstly, we validated their ability to form secondary structures by circular dichroism (CD) spectroscopy, a standard technique for G4 structure analysis (59). DNA G4 components with varying inhibitory activities (see Fig 2) were analyzed to reveal their structural conformations, namely DNA13, DNA15, DNA30 and DNA40. Henceforth, all synthetic G4 components will be named according to the nucleic acid level at which they operate and the relative protein expression units (REU) they encode, e.g. DNA.70REU (DNA13), DNA.50REU (DNA15), DNA.30REU (DNA30) and DNA.10REU (DNA40; see Supplementary Table 1 for all G4 nucleic acid sequences and names). As shown in Figure 3A, all four components produced classical distinctive G4 CD spectra (60), where the mutated control sequence did not show characteristic G4 peaks and troughs (Fig. 3A). We note that G4s with highest inhibitory activities (DNA.10REU and DNA.30REU) formed parallel structures (positive peak at 260 nm, negative peak at 240 nm), while medium (DNA.50REU) and low (DNA.70REU) strength components adopted hybrid and anti-parallel (positive peaks at 295 and 260 nm, negative peak at 240 nm) conformations respectively (61); the mutated sequence displayed the spectrum typical of unstructured single-stranded DNA (62). Based on these data, and our previous finding that regulatory activities were correlated with sequence feature compositions, we concluded that synthetic components’ relative performances were underpinned by their varying propensities to form and maintain G4 structures. Although we note that some *in vivo* applications, such as gene therapy, may require a more detailed analysis of intracellular secondary structure formation dynamics using G4-specific fluorescent probes (63) or antibodies (64).

**Figure 3.**
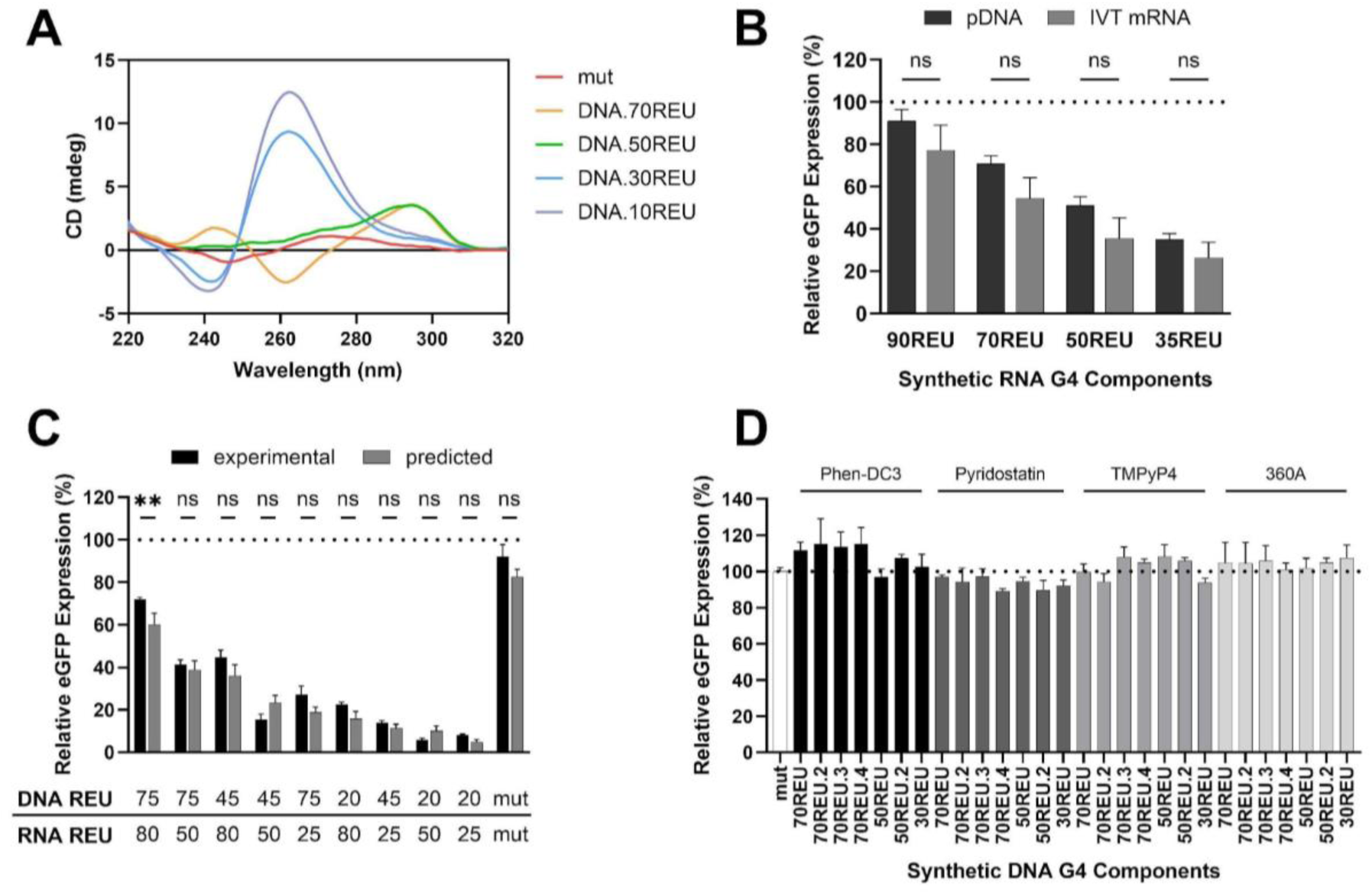
**A)** Circular dichroism spectra of DNA oligonucleotides containing varying G-quadruplex (G4) motifs (all G4 motif sequences listed in Supplementary Table 1). CHO cells were transiently transfected with **B)** Synthetic eGFP-encoding mRNA molecules harboring different RNA G4 components, and **C)** eGFP reporter plasmids containing varying combinations of DNA and RNA G4 elements. eGFP levels were quantified 48 h post-transfection and compared to those driven by b) the same G4s in a DNA plasmid context, and c) predicted values, assuming constituent G4 pairs act synergistically (see Fig 2). **D)** CHO cells were stably transfected with G4-reporter plasmids coexpressing a puromycin selection marker gene. Stable cell pools were created and subsequently exposed to varying chemical ligands for 48 h prior to eGFP quantification. In B and C data are expressed as a percentage of the production exhibited by the control CMV-IE1 proximal promoter-BPCU (ΔG4) construct (dotted line). Values represent the mean + SD of three independent experiments (n = 3, each performed in triplicate). In D, data are expressed as a percentage of the production achieved in control media (Δligand, containing DMSO or water), normalized to the effect of ligand addition on G4 mutant (white bar). Values represent the mean + SD of three independent experiments (n = 3, each performed in duplicate).

Given the dramatic rise in applications utilizing synthetic mRNA, including protein replacement therapy, cancer immunotherapy and vaccines against infectious diseases (4, 65), we next sought to validate that our RNA components display predictable performance when employed directly in mRNA molecular contexts. eGFP-mRNA molecules harboring RNA elements of varying inhibitory strengths were produced by *in vitro* transcription reactions, where purified transcripts were identical to those created intracellularly from DNA reporter plasmids (i.e. comprising the AZ 5’ UTR containing a single G4 element in position +6, upstream of the eGFP CDS; see Fig 1). Synthetic mRNAs were transfected into CHO cells prior to quantification of eGFP protein levels after 48 h. These data show that synthetic element performance in DNA vector and mRNA transcript contexts is highly correlated (Pearson’s correlation coefficient, r = 0.99, p = 0.008) where RNA.90REU, RNA.70REU, RNA.50REU and RNA35.REU drove eGFP expression levels of 75%, 55%, 35% and 25% respectively, relative to the unengineered control (Fig 3B). Accordingly, synthetic RNA G4 components from the library can be utilized predictably to precisely control recombinant protein production in both plasmid DNA (e.g. biopharmaceutical production) and synthetic mRNA (e.g. cancer vaccines) molecular formats. We note that we were unable to synthesize mRNAs containing element RNA.5REU, presumably due to secondary structure-based effects, and therefore use of very strong G4s (i.e. REUs < 20) may require optimization of mRNA manufacturing processes.

Synthetic mammalian transcriptional and translational control elements are typically developed in isolation, where a lack of standardization, such as the use of varying genetic architectures and test systems, complicates efforts to use them in combination (66, 67). G4-based regulatory controllers operating within a standardized BPCU should permit predictable, simultaneous titration of transcription and translation rates via selection of appropriate DNA and RNA parts. To test this, we created constructs containing all possible combinations of exemplar high, medium and low strength DNA (DNA.20REU, DNA.45REU, DNA.75REU) and RNA (RNA.25REU, RNA.50REU, RNA.80REU) components. As shown in Figure 3C, dual-controllers drove eGFP expression levels in CHO cells that were highly correlated with predicted values (r = 0.96, p = < 0.0001), assuming synergistic effects of constituent elements. Indeed, all observed activities were within ± 12 REU of expected outputs, indicating that composite parts functioned as expected to reduce the rate of both recombinant gene transcription and recombinant mRNA translation. We therefore concluded that DNA and RNA elements from the library can be predictably combined to achieve user-defined, stringent two-level control of recombinant protein expression, which may be particularly advantageous for applications that necessitate product biosynthesis rates to be maintained within defined limits, such as manufacture of toxic molecules.

Finally, we evaluated the potential to convert our G4-based controllers into inducible expression systems. Many endogenous G4 sequences interact with chemical ligands that can destabilize or stabilize their structure to increase (ON-systems) or decrease (OFF-systems) protein expression levels (68, 69). Such genetic switches facilitate sophisticated control strategies, including the ability to enact data- and process-responsive titration of system outputs (70, 71). We tested four well-known G4 ligands (Phen-DC3, Pyridostatin, TMPyP4, and 360A (69, 72)) in combination with seven distinct DNA elements that were selected to i) have varying feature compositions, based on findings that ligands may have specific binding preferences (73, 74), and ii) exhibit medium to low strength (30 – 70 REU), to permit observation of up- and down-regulatory effects. These 28 expression systems were evaluated in an industrially-relevant context, namely stable protein production in CHO cells, where there is a design objective to switch expression ‘ON’ once maximum cell densities are reached in order to decouple growth and production phases during biomanufacturing processes (75). As shown in Figure 3D, eGFP expression levels driven by synthetic G4 components (DNA.30REU, DNA.50REU, DNA.50REU.2, DNA.70REU, DNA.70REU.2 DNA.70REU.3 and DNA.70REU.4) were not significantly affected by any of the tested ligands. Further tests demonstrated that RNA G4 component activity was similarly unchanged by addition of these chemicals (data not shown). Failure to create functional ON and OFF switches may be due to ligand’s inabilities to interact with unnatural synthetic G4 sequences. Alternatively, maximum useable chemical concentrations (not reducing cell viability by > 10%) may have been insufficient to significantly regulate the artificially large number of G4 motifs present per cell, where stably transfected CHO cells typically contain a high number of integrated gene copies (76, 77). These findings, coupled with broad-acting ligands potentially causing off-target effects on host cell function by binding to multiple endogenous structures, indicate that development of inducible systems for bioindustrial contexts will require design of bespoke, highly specific synthetic G4 modulators (74, 78). Such genetic switches would also have potential use in gene therapy by facilitating dynamic titration of therapeutic protein levels in response to patient data profiles.

### 3.4 Validating synthetic G4 component performance for bioindustrial applications

As G4 controllers function via interference with the general cellular biosynthetic machinery, rather than site-specific regulators such as transcription factors, they should exhibit modular performances where activities are not affected by the chosen mammalian cell host. Indeed, the ability to predictably deploy these components in new bioindustrial cell contexts is a potential key advantage of this technology, particularly for *in vivo* therapeutic applications which may target any one of hundreds of distinct human cell types (79, 80). To test this, we profiled the performance of our G4 elements in two additional bioindustrially-relevant cellular environments, namely i) Human Embryonic Kidney (HEK) cells, a commonly utilized biomanufacturing host for recombinant protein and viral vector production (81), and ii) HepG2 cells, a model cell line representing a common target tissue for gene therapy (82, 83). Reporter plasmids harboring DNA and RNA G4 components of varying activities were individually transfected into each cell type and eGFP expression was quantified 48 h later. As illustrated in Figure 4A, DNA and RNA components performed as expected in both new cell contexts, each driving five distinct predictable levels of protein production ranging from very low to high. Indeed, previously observed activities in CHO cells were highly correlated with those seen in HEK (Pearson’s correlation coefficient, r = 0.93, p = < 0.0001) and HepG2 (r = 0.97, p = < 0.0001). We note that although discrete encoded protein production levels were maintained, construct inhibitory activities were generally slightly weaker in these cell types, as compared to CHO cells, resulting in higher mean expression outputs in both HEK (+7 REU) and HepG2 (+17 REU). This aligns with previous studies showing relative G4 regulatory activities are maintained between cell lines but overall suppressive effects are somewhat different (41), potentially due to varying ion compositions (84, 85). Nevertheless, our findings confirm that these synthetic G4s can be predictably utilized as ‘off-the-shelf’ components to optimally titrate recombinant protein expression levels in new cell types of interest.

**Figure 4.**
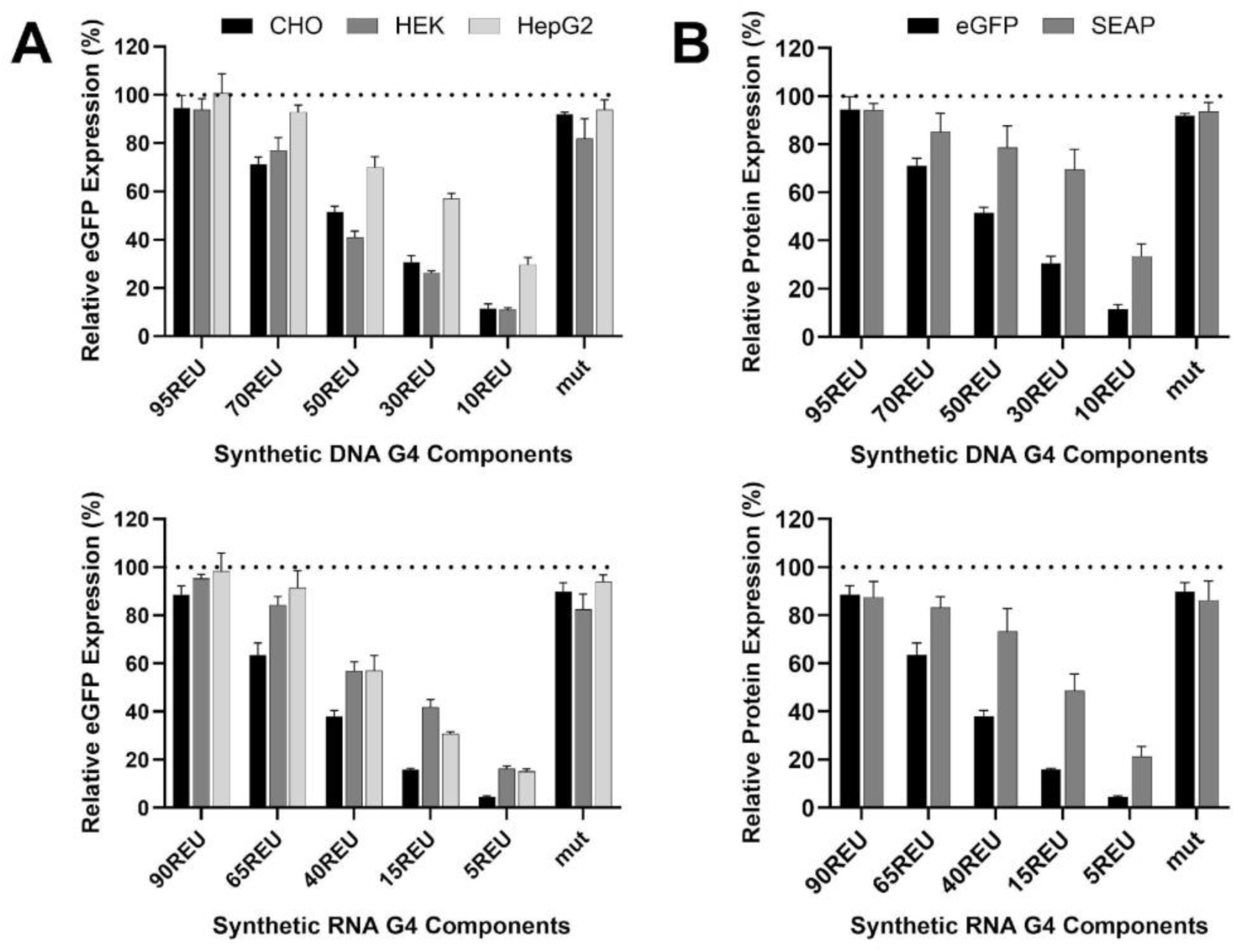
**A)** eGFP-reporter plasmids containing DNA or RNA G4-components with varying activities were transfected into CHO, HEK and HepG2 cells, prior to protein quantification 48 h later. **B)** CHO cells were transfected with reporter plasmids encoding eGFP or SEAP production under the control of varying-strength DNA or RNA G4 elements. Data are expressed as a percentage of the production exhibited by the control CMV-IE1 proximal promoter-BPCU (ΔG4) construct (dotted line). Values represent the mean + SD of three independent experiments (n = 3, each performed in triplicate).

We next evaluated the ability to utilize synthetic G4 constructs to routinely tailor expression of new target recombinant proteins, using Secreted Alkaline Phosphatase (SEAP) as an exemplar secreted, glycosylated molecule. Reporter vectors encoding SEAP expression driven by G4 controllers of varying strength were transfected into CHO cells prior to product quantification after 48 h. SEAP outputs from G4 elements were highly correlated with expected values (r = 0.91, p = 0.0002) although product expression levels, relative to the standard BPCU, were higher than those observed for eGFP (Fig 4B). The latter observation is likely due to product-specific biosynthetic processing kinetics (i.e. impact of post-translational processing steps), where relative mature protein levels do not directly correlate with relative mRNA transcript and nascent polypeptide quantities (86–88). Accordingly, where very low protein levels are required (i.e. < 10% of that driven by the standard expression cassette lacking a G4 element), we recommend creating vector-optimization spaces that include dual-controllers encoding minimal expression outputs (e.g. DNA.10REU and RNA.5REU utilized in combination, see Fig 3C). Nevertheless, the selected G4 component mini library performed as designed by facilitating six discrete expression set-points for a new protein-of-interest (i.e. 20, 30, 50, 70, 85, and 100 relative protein expression units), validating their product-agnostic functionality.

Finally, we incorporated our G4 controller technology into a current high-value bioindustrial process, namely cell line development for protein biopharmaceutical production which involves derivation of optimized CHO cell host, DNA expression vector combinations. The dominant class of biopharmaceuticals, monoclonal antibodies (mAb), consist of separate heavy chain (HC) and light chain (LC) polypeptides, where it is well known that the optimal HC:LC expression ratio is highly product specific, ranging from 1:1 to 1:10 depending on antibody folding and assembly dynamics (89–91). We utilized four DNA G4 components of varying strength, DNA.REU70, DNA.REU50, DNA.REU30 and DNA.REU10, alongside the unengineered BPCU construct (i.e. 100 REU) to independently control LC and HC expression for an exemplar AZ mAb (mAbT; Fig 5A). All possible component combinations were tested in transient fed-batch production processes, where the 25 unique encoded HC:LC expression ratios facilitated a wide range of mAbT titers (Fig 5B). As illustrated in Figure 5C, rapidly testing a standardized vector solution space revealed that i) an encoded HC:LC expression ratio of ∼1:2 enabled highest product titer, ii) mAbT yields were positively correlated with increasing LC quantities, and iii) HC levels needed to be precisely controlled to achieve an optimal set-point, presumably due to excess intracellular HC causing adverse effects on cell factory performance (89, 92). We therefore concluded that a G4 element-based screening platform could derive optimal expression vector designs for new proteins, and could also be utilized as a diagnostic tool to interrogate the cell’s relative ability to synthesize a given polypeptide, guiding product-specific cell engineering solutions (93–95). We anticipate that this technology will be particularly useful for the development of difficult-to-express proteins, where suboptimal polypeptide biosynthesis kinetics dramatically restricts product yield and quality, and next-generation molecules that require expression of ≥ 3 composite chains to be precisely stoichiometrically balanced (96–98).

**Figure 5.**
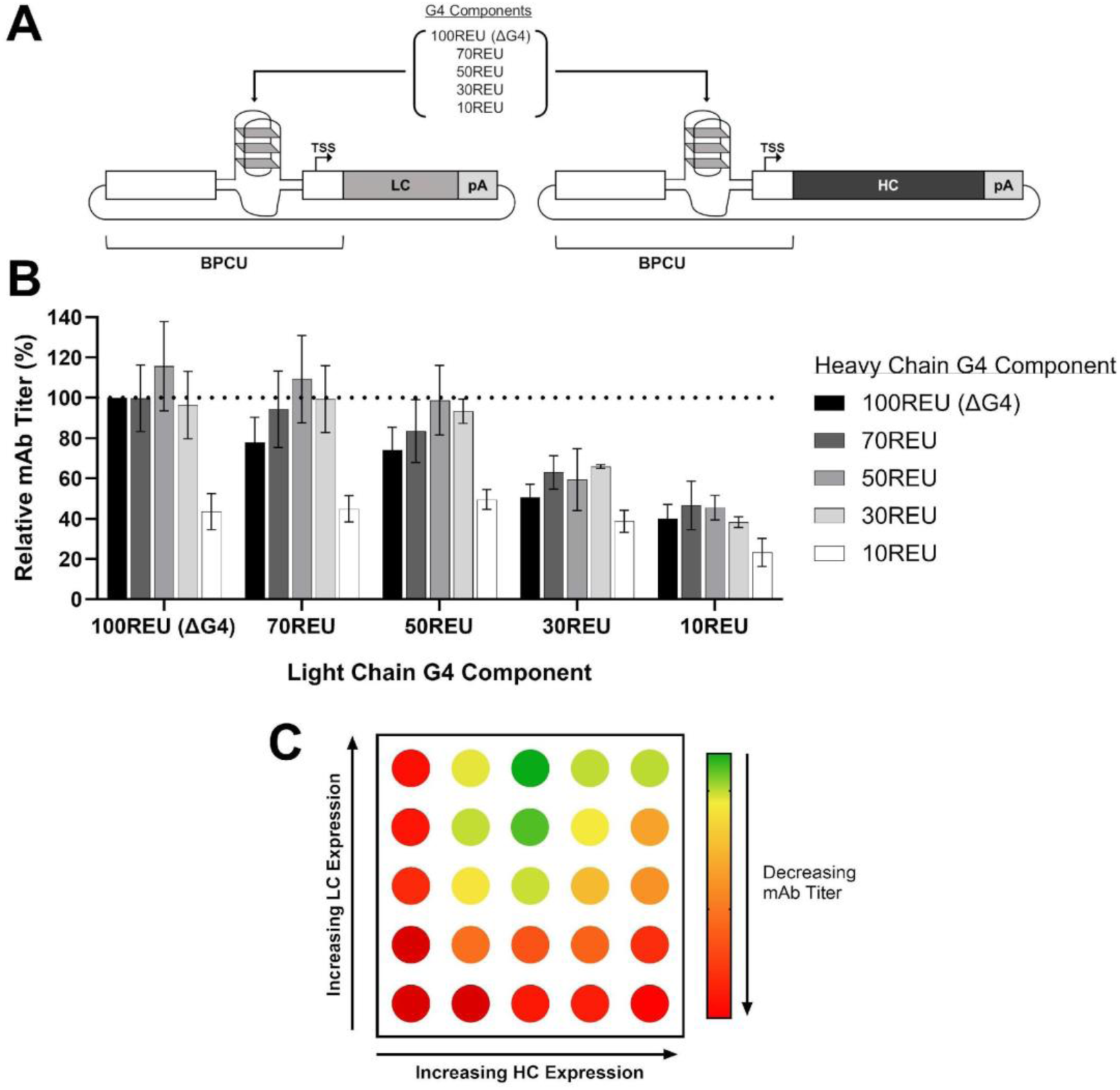
**A)** Vector design platform to optimize polypeptide expression ratios for multichain biopharmaceutical products. **B)** CHO cells were co-transfected with separate plasmids (at an equimolar ratio) encoding mAbT heavy (HC) and light (LC) chain expression under the control of varying-strength G4 components (sequences listed in Supplementary Table 1). Cells were incubated for 5 days, including addition of feed 48 h post-transfection, prior to quantification of mAb titer. Data are shown as a percentage of the production achieved using the unengineered BPCU (ΔG4) construct to drive expression of both HC and LC (dotted line). Values represent the mean + SD of three independent experiments (n = 3, each performed in triplicate). **C)** Vector solution space identifying mAbT product-specific design rules.

## Conclusion

The synthetic G4 components presented in this study can be used predictably to precisely tune mammalian recombinant protein expression levels. Specifically created to function within a bioindustry-compatible genetic chassis, these elements can provide simple off-the-shelf solutions to current protein production challenges. Shown to reliably encode a wide-range of discrete protein biosynthesis set-points in both CHO and HEK cells, they can be harnessed to i) optimize polypeptide expression ratios for difficult-to-express multi-chain molecules, such as tri-specific antibodies, and Adeno-associated viral vectors, and ii) enact complex multigene cell factory engineering strategies (5, 98, 99). Having demonstrated the technology’s performance for transient production contexts (100), we note that implementation in stable expression systems will require comprehensive validation in relevant industrial manufacturing platforms (e.g. using standard genome integration methodologies, clonal selection procedures, etc. (101)).

Demonstrated to titrate protein expression rates predictably in varying molecular formats (i.e. mRNA and DNA) and mammalian cell types, synthetic G4s could be deployed in gene therapy applications to coordinate therapeutic efficacy with product safety by driving optimized expression levels that achieve intended medical outcomes without causing off-target effects on cell behavior (102). Moreover, for the growing number of products that target multigenic disorders, they could be used to precisely balance the intracellular levels of multiple therapeutic proteins in order to achieve required, physiologically-relevant expression ratios (103). Given that in *in vivo* applications, genetic regulatory elements become ‘part of the product’ introduced into patients, their successful incorporation into such therapeutics will require extensive testing in animal models.

As they enable simultaneous precision control of both transcription and translation rates, G4 components can also be utilized in combination with other gene expression modulation technologies including cell-type specific promoters (11) and exogenous additives that act as ON or OFF switches (104). Indeed, the ability to enact two-level protein expression control with a single motif (i.e. using the same G4 sequence in the core promoter and 5’ UTR) provides an ideal basis for constructing non-leaky inducible expression systems that can coordinate product biosynthesis kinetics with bioreactor processes, cell line development workflows and patient requirements (75, 105, 106). However, this will require the design of cognate regulatory-compliant chemical ligands that exhibit highly G4-specific stabilization or destabilization activities without inducing off-target effects that disrupt other required host cell functionalities (107). In conclusion, the synthetic genetic elements introduced in this study provide a new toolkit to precisely tailor mammalian recombinant protein expression in high-value bioindustrial applications, where future work will focus on the development of next-generation small molecule-inducible ON and OFF switches.

## References

1. Brown, A.J., Gibson, S.J., Hatton, D., Arnall, C.L. and James, D.C. (2019) Whole synthetic pathway engineering of recombinant protein production. Biotechnol. Bioeng., 116, 375–387.

2. Wang, T.Y. and Guo, X. (2020) Expression vector cassette engineering for recombinant therapeutic production in mammalian cell systems. Appl. Microbiol. Biotechnol., 104, 5673–5688.

3. Poletti, V. and Mavilio, F. (2021) Designing lentiviral vectors for gene therapy of genetic diseases. Viruses, 13.

4. Qin, S., Tang, X., Chen, Y., Chen, K., Fan, N., Xiao, W., Zheng, Q., Li, G., Teng, Y., Wu, M., et al. (2022) mRNA-based therapeutics: powerful and versatile tools to combat diseases. Signal Transduct. Target. Ther., 7, 1–35.

5. Eisenhut, P., Marx, N., Borsi, G., Papež, M., Ruggeri, C., Baumann, M. and Borth, N. (2024) Manipulating gene expression levels in mammalian cell factories: An outline of synthetic molecular toolboxes to achieve multiplexed control. N. Biotechnol., 79, 1–19.

6. Cring, M.R. and Sheffield, V.C. (2022) Gene therapy and gene correction: targets, progress, and challenges for treating human diseases. Gene Ther., 29, 3–12.

7. Sou, S.N., Harris, C.L., Williams, R., Kozub, D., Zurlo, F., Patel, Y.D., Kallamvalli Illam Sankaran, P., Daramola, O., Brown, A., James, D.C., et al. (2023) CHO synthetic promoters improve expression and product quality of biotherapeutic proteins. Biotechnol. Prog., 39, e3348.

8. Li, Z.M., Fan, Z.L., Wang, X.Y. and Wang, T.Y. (2022) Factors Affecting the Expression of Recombinant Protein and Improvement Strategies in Chinese Hamster Ovary Cells. Front. Bioeng. Biotechnol., 10.

9. Samoudi, M., Masson, H.O., Kuo, C.C., Robinson, C.M. and Lewis, N.E. (2021) From omics to cellular mechanisms in mammalian cell factory development. Curr. Opin. Chem. Eng., 32, 100688.

10. O’Neill, P., Mistry, R.K., Brown, A.J. and James, D.C. (2023) Protein-Specific Signal Peptides for Mammalian Vector Engineering. ACS Synth. Biol., 12, 2339–2352.

11. Johnson, A.O., Fowler, S.B., Webster, C.I., Brown, A.J. and James, D.C. (2022) Bioinformatic Design of Dendritic Cell-Specific Synthetic Promoters. ACS Synth. Biol., 11, 1613–1626.

12. Cao, J., Novoa, E.M., Zhang, Z., Chen, W.C.W., Liu, D., Choi, G.C.G., Wong, A.S.L., Wehrspaun, C., Kellis, M. and Lu, T.K. (2021) High-throughput 5′ UTR engineering for enhanced protein production in non-viral gene therapies. Nat. Commun., 12, 1–10.

13. Johari, Y.B., Mercer, A.C., Liu, Y., Brown, A.J. and James, D.C. (2021) Design of synthetic promoters for controlled expression of therapeutic genes in retinal pigment epithelial cells. Biotechnol. Bioeng., 118, 2001–2015.

14. Skopenkova, V. V., Egorova, T. V. and Bardina, M. V. (2021) Muscle-Specific Promoters for Gene Therapy. Acta Naturae, 13, 47–58.

15. McGraw, C.E., Peng, D. and Sandoval, N.R. (2020) Synthetic biology approaches: the next tools for improved protein production from CHO cells. Curr. Opin. Chem. Eng., 30, 26–33.

16. Leppek, K., Das, R. and Barna, M. (2018) Functional 5′ UTR mRNA structures in eukaryotic translation regulation and how to find them. Nat. Rev. Mol. Cell Biol., 19, 158–174.

17. Eisenhut, P., Mebrahtu, A., Moradi Barzadd, M., Thalén, N., Klanert, G., Weinguny, M., Sandegren, A., Su, C., Hatton, D., Borth, N., et al. (2020) Systematic use of synthetic 5’-UTR RNA structures to tune protein translation improves yield and quality of complex proteins in mammalian cell factories. Nucleic Acids Res., 48, e119–e119.

18. Sen, D. and Gilbert, W. (1988) Formation of parallel four-stranded complexes by guanine-rich motifs in DNA and its implications for meiosis. Nature, 334, 364–366.

19. Chambers, V.S., Marsico, G., Boutell, J.M., Di Antonio, M., Smith, G.P. and Balasubramanian, S. (2015) High-throughput sequencing of DNA G-quadruplex structures in the human genome. Nat. Biotechnol., 33, 877–881.

20. Kwok, C.K., Marsico, G., Sahakyan, A.B., Chambers, V.S. and Balasubramanian, S. (2016) RG4-seq reveals widespread formation of G-quadruplex structures in the human transcriptome. Nat. Methods, 13, 841–844.

21. Masai, H. and Tanaka, T. (2020) G-quadruplex DNA and RNA: Their roles in regulation of DNA replication and other biological functions. Biochem. Biophys. Res. Commun., 531, 25–38.

22. Lago, S., Nadai, M., Cernilogar, F.M., Kazerani, M., Domíniguez Moreno, H., Schotta, G. and Richter, S.N. (2021) Promoter G-quadruplexes and transcription factors cooperate to shape the cell type-specific transcriptome. Nat. Commun., 12, 1–13.

23. Robinson, J., Raguseo, F., Nuccio, S.P., Liano, D. and Di Antonio, M. (2021) DNA G-quadruplex structures: More than simple roadblocks to transcription? Nucleic Acids Res., 49, 8419–8431.

24. Varshney, D., Spiegel, J., Zyner, K., Tannahill, D. and Balasubramanian, S. (2020) The regulation and functions of DNA and RNA G-quadruplexes. Nat. Rev. Mol. Cell Biol., 21, 459–474.

25. Song, J., Perreault, J.-P., Topisirovic, I. and Richard, S. (2016) RNA G-quadruplexes and their potential regulatory roles in translation. Translation, 4, e1244031.

26. Bochman, M.L., Paeschke, K. and Zakian, V.A. (2012) DNA secondary structures: Stability and function of G-quadruplex structures. Nat. Rev. Genet., 13, 770–780.

27. Wang, W., Hu, S., Gu, Y., Yan, Y., Stovall, D.B., Li, D. and Sui, G. (2020) Human MYC G-quadruplex: From discovery to a cancer therapeutic target. Biochim. Biophys. Acta - Rev. Cancer, 1874, 188410.

28. Morgan, R.K., Batra, H., Gaerig, V.C., Hockings, J. and Brooks, T.A. (2016) Identification and characterization of a new G-quadruplex forming region within the kRAS promoter as a transcriptional regulator. Biochim. Biophys. Acta - Gene Regul. Mech., 1859, 235–245.

29. Kumari, S., Bugaut, A. and Balasubramanian, S. (2008) Position and stability are determining factors for translation repression by an RNA G-quadruplex-forming sequence within the 5′ UTR of the NRAS proto-oncogene. Biochemistry, 47, 12664–12669.

30. Yan, J., Zhao, D., Dong, L., Pan, S., Hao, F. and Guan, Y. (2017) A novel G-quadruplex motif in the Human MET promoter region. Biosci. Rep., 37.

31. Onel, B., Carver, M., Wu, G., Timonina, D., Kalarn, S., Larriva, M. and Yang, D. (2016) A New G-Quadruplex with Hairpin Loop Immediately Upstream of the Human BCL2 P1 Promoter Modulates Transcription. J. Am. Chem. Soc., 138, 2563–2570.

32. Shahid, R., Bugaut, A. and Balasubramanian, S. (2010) The BCL-2 5′ untranslated region contains an RNA G-quadruplex-forming motif that modulates protein expression. Biochemistry, 49, 8300–8306.

33. Huang, W., Smaldino, P.J., Zhang, Q., Miller, L.D., Cao, P., Stadelman, K., Wan, M., Giri, B., Lei, M., Nagamine, Y., et al. (2012) Yin Yang 1 contains G-quadruplex structures in its promoter and 5′-UTR and its expression is modulated by G4 resolvase 1. Nucleic Acids Res., 40, 1033–1049.

34. Romanova, N. and Noll, T. (2018) Engineered and Natural Promoters and Chromatin-Modifying Elements for Recombinant Protein Expression in CHO Cells. Biotechnol. J., 13, 1700232.

35. Brown, A.J. and James, D.C. (2016) Precision control of recombinant gene transcription for CHO cell synthetic biology. Biotechnol. Adv., 34, 492–503.

36. Patel, Y.D., Brown, A.J., Zhu, J., Rosignoli, G., Gibson, S.J., Hatton, D. and James, D.C. (2021) Control of Multigene Expression Stoichiometry in Mammalian Cells Using Synthetic Promoters. ACS Synth. Biol., 10, 1155–1165.

37. Williams, D.J., Puhl, H.L. and Ikeda, S.R. (2010) A simple, highly efficient method for heterologous expression in mammalian primary neurons using cationic lipid-mediated mRNA transfection. Front. Neurosci., 4, 1761.

38. Brown, A.J., Sweeney, B., Mainwaring, D.O. and James, D.C. (2014) Synthetic promoters for CHO cell engineering. Biotechnol. Bioeng., 111, 1638–1647.

39. Johari, Y.B., Scarrott, J.M., Pohle, T.H., Liu, P., Mayer, A., Brown, A.J. and James, D.C. (2022) Engineering of the CMV promoter for controlled expression of recombinant genes in HEK293 cells. Biotechnol. J., 17, 2200062.

40. Au, H.K.E., Isalan, M. and Mielcarek, M. (2022) Gene Therapy Advances: A Meta-Analysis of AAV Usage in Clinical Settings. Front. Med., 8, 809118.

41. Halder, K., Wieland, M. and Hartig, J.S. (2009) Predictable suppression of gene expression by 5′-UTR-based RNA quadruplexes. Nucleic Acids Res., 37, 6811–6817.

42. Xia, W., Bringmann, P., McClary, J., Jones, P.P., Manzana, W., Zhu, Y., Wang, S., Liu, Y., Harvey, S., Madlansacay, M.R., et al. (2006) High levels of protein expression using different mammalian CMV promoters in several cell lines. Protein Expr. Purif., 45, 115–124.

43. Qin, J.Y., Zhang, L., Clift, K.L., Hulur, I., Xiang, A.P., Ren, B.Z. and Lahn, B.T. (2010) Systematic comparison of constitutive promoters and the doxycycline-inducible promoter. PLoS One, 5, 10611.

44. Juven-Gershon, T. and Kadonaga, J.T. (2010) Regulation of gene expression via the core promoter and the basal transcriptional machinery. Dev. Biol., 339, 225–229.

45. Kozak, M. (1989) Circumstances and Mechanisms of Inhibition of Translation by Secondary Structure in Eucaryotic mRNAs. Mol. Cell. Biol., 9, 5134–5142.

46. Huppert, J.L. and Balasubramanian, S. (2007) G-quadruplexes in promoters throughout the human genome. Nucleic Acids Res., 35, 406–413.

47. Beaudoin, J.D. and Perreault, J.P. (2010) 5′-UTR G-quadruplex structures acting as translational repressors. Nucleic Acids Res., 38, 7022–7036.

48. Beaudoin, J.D., Jodoin, R. and Perreault, J.P. (2014) New scoring system to identify RNA G-quadruplex folding. Nucleic Acids Res., 42, 1209–1223.

49. Kikin, O., D’Antonio, L. and Bagga, P.S. (2006) QGRS Mapper: A web-based server for predicting G-quadruplexes in nucleotide sequences. Nucleic Acids Res., 34, W676–W682.

50. Agarwal, T., Roy, S., Kumar, S., Chakraborty, T.K. and Maiti, S. (2014) In the sense of transcription regulation by G-quadruplexes: Asymmetric effects in sense and antisense strands. Biochemistry, 53, 3711–3718.

51. Patwardhan, R.P., Lee, C., Litvin, O., Young, D.L., Pe’Er, D. and Shendure, J. (2009) High-resolution analysis of DNA regulatory elements by synthetic saturation mutagenesis. Nat. Biotechnol., 27, 1173–1175.

52. Pandey, S., Agarwala, P. and Maiti, S. (2013) Effect of loops and G-quartets on the stability of RNA G-quadruplexes. J. Phys. Chem. B, 117, 6896–6905.

53. Zhang, A.Y.Q., Bugaut, A. and Balasubramanian, S. (2011) A sequence-independent analysis of the loop length dependence of intramolecular RNA G-quadruplex stability and topology. Biochemistry, 50, 7251–7258.

54. Petraccone, L., Erra, E., Duro, I., Esposito, V., Randazzo, A., Mayol, L., Mattia, C.A., Barone, G. and Giancola, C. (2005) Relative stability of quadruplexes containing different number of G-tetrads. In Nucleosides, Nucleotides and Nucleic Acids. Vol. 24, pp. 757–760.

55. Yu, Y.Z., Ma, Y., Xu, W.H., Wang, S. and Sun, Z.W. (2015) Combinations of various CpG motifs cloned into plasmid backbone modulate and enhance protective immunity of viral replicon DNA anthrax vaccines. Med. Microbiol. Immunol., 204, 481–491.

56. Chan, Y.K., Wang, S.K., Chu, C.J., Copland, D.A., Letizia, A.J., Verdera, H.C., Chiang, J.J., Sethi, M., Wang, M.K., Neidermyer, W.J., et al. (2021) Engineering adeno-associated viral vectors to evade innate immune and inflammatory responses. Sci. Transl. Med., 13, 3438.

57. Hoose, A., Vellacott, R., Storch, M., Freemont, P.S. and Ryadnov, M.G. (2023) DNA synthesis technologies to close the gene writing gap. Nat. Rev. Chem., 7, 144–161.

58. Fath, S., Bauer, A.P., Liss, M., Spriestersbach, A., Maertens, B., Hahn, P., Ludwig, C., Schäfer, F., Graf, M. and Wagner, R. (2011) Multiparameter RNA and codon optimization: A standardized tool to assess and enhance autologous mammalian gene expression. PLoS One, 6, e17596.

59. Grün, J.T. and Schwalbe, H. (2022) Folding dynamics of polymorphic G-quadruplex structures. Biopolymers, 113, e23477.

60. del Villar-Guerra, R., Trent, J.O. and Chaires, J.B. (2018) G-Quadruplex Secondary Structure Obtained from Circular Dichroism Spectroscopy. Angew. Chemie - Int. Ed., 57, 7171–7175.

61. Kejnovská, I., Renčiuk, D., Palacký, J. and Vorlíčková, M. (2019) CD Study of the G-Quadruplex Conformation. In Methods in Molecular Biology. Humana Press Inc., Vol. 2035, pp. 25–44.

62. Johnson, W.C. (1996) Determination of the Conformation of Nucleic Acids by Electronic CD. In Circular Dichroism and the Conformational Analysis of Biomolecules. Springer, Boston, MA, pp. 433–468.

63. Di Antonio, M., Ponjavic, A., Radzevičius, A., Ranasinghe, R.T., Catalano, M., Zhang, X., Shen, J., Needham, L.M., Lee, S.F., Klenerman, D., et al. (2020) Single-molecule visualization of DNA G-quadruplex formation in live cells. Nat. Chem., 12, 832–837.

64. Biffi, G., Tannahill, D., McCafferty, J. and Balasubramanian, S. (2013) Quantitative visualization of DNA G-quadruplex structures in human cells. Nat. Chem., 5, 182–186.

65. Vavilis, T., Stamoula, E., Ainatzoglou, A., Sachinidis, A., Lamprinou, M., Dardalas, I. and Vizirianakis, I.S. (2023) mRNA in the Context of Protein Replacement Therapy. Pharmaceutics, 15.

66. Sample, P.J., Wang, B., Reid, D.W., Presnyak, V., McFadyen, I.J., Morris, D.R. and Seelig, G. (2019) Human 5′ UTR design and variant effect prediction from a massively parallel translation assay. Nat. Biotechnol., 37, 803–809.

67. Cazier, A.P. and Blazeck, J. (2021) Advances in promoter engineering: Novel applications and predefined transcriptional control. Biotechnol. J., 16, 2100239.

68. Chaudhary, S., Kumar, M. and Kaushik, M. (2022) Interface of G-quadruplex with both stabilizing and destabilizing ligands for targeting various diseases. Int. J. Biol. Macromol., 219, 414–427.

69. Yan, M.P., Wee, C.E., Yen, K.P., Stevens, A. and Wai, L.K. (2023) G-quadruplex ligands as therapeutic agents against cancer, neurological disorders and viral infections. Future Med. Chem., 15, 1987–2009.

70. Pardi, M.L., Wu, J., Kawasaki, S. and Saito, H. (2022) Synthetic RNA-based post-transcriptional expression control methods and genetic circuits. Adv. Drug Deliv. Rev., 184, 114196.

71. Verbič, A., Praznik, A. and Jerala, R. (2021) A guide to the design of synthetic gene networks in mammalian cells. FEBS J., 288, 5265–5288.

72. Figueiredo, J., Mergny, J.L. and Cruz, C. (2024) G-quadruplex ligands in cancer therapy: Progress, challenges, and clinical perspectives. Life Sci., 340, 122481.

73. Wu, G., Tillo, D., Ray, S., Chang, T.C., Schneekloth, J.S., Vinson, C. and Yang, D. (2020) Custom G4 microarrays reveal selective G-quadruplex recognition of small molecule BMVC: A large-scale assessment of ligand binding selectivity. Molecules, 25, 3465.

74. Duarte, A.R., Cadoni, E., Ressurreição, A.S., Moreira, R. and Paulo, A. (2018) Design of Modular G-quadruplex Ligands. ChemMedChem, 13, 869–893.

75. Donaldson, J.S., Dale, M.P. and Rosser, S.J. (2021) Decoupling Growth and Protein Production in CHO Cells: A Targeted Approach. Front. Bioeng. Biotechnol., 9, 349.

76. Lee, Z., Wan, J., Shen, A. and Barnard, G. (2024) Gene copy number, gene configuration and LC/HC mRNA ratio impact on antibody productivity and product quality in targeted integration CHO cell lines. Biotechnol. Prog., 40, e3433.

77. Joubert, S., Stuible, M., Lord-Dufour, S., Lamoureux, L., Vaillancourt, F., Perret, S., Ouimet, M., Pelletier, A., Bisson, L., Mahimkar, R., et al. (2023) A CHO stable pool production platform for rapid clinical development of trimeric SARS-CoV-2 spike subunit vaccine antigens. Biotechnol. Bioeng., 120, 1746–1761.

78. Berner, A., Das, R.N., Bhuma, N., Golebiewska, J., Abrahamsson, A., Andréasson, M., Chaudhari, N., Doimo, M., Bose, P.P., Chand, K., et al. (2024) G4-Ligand-Conjugated Oligonucleotides Mediate Selective Binding and Stabilization of Individual G4 DNA Structures. J. Am. Chem. Soc., 146, 6926–6935.

79. Regev, A., Teichmann, S.A., Lander, E.S., Amit, I., Benoist, C., Birney, E., Bodenmiller, B., Campbell, P., Carninci, P., Clatworthy, M., et al. (2017) The human cell atlas. Elife, 6.

80. Shchaslyvyi, A.Y., Antonenko, S. V., Tesliuk, M.G. and Telegeev, G.D. (2023) Current State of Human Gene Therapy: Approved Products and Vectors. Pharmaceuticals, 16, 1416.

81. Tan, E., Chin, C.S.H., Lim, Z.F.S. and Ng, S.K. (2021) HEK293 Cell Line as a Platform to Produce Recombinant Proteins and Viral Vectors. Front. Bioeng. Biotechnol., 9.

82. Arzumanian, V.A., Kiseleva, O.I. and Poverennaya, E. V. (2021) The curious case of the HepG2 cell line: 40 years of expertise. Int. J. Mol. Sci., 22, 13135.

83. Zabaleta, N., Unzu, C., Weber, N.D. and Gonzalez-Aseguinolaza, G. (2023) Gene therapy for liver diseases — progress and challenges. Nat. Rev. Gastroenterol. Hepatol., 20, 288–305.

84. Bhattacharyya, D., Arachchilage, G.M. and Basu, S. (2016) Metal cations in G-quadruplex folding and stability. Front. Chem., 4, 38.

85. Zhang, Z.H., Qian, S.H., Wei, D. and Chen, Z.X. (2023) In vivo dynamics and regulation of DNA G-quadruplex structures in mammals. Cell Biosci., 13.

86. Mathias, S., Wippermann, A., Raab, N., Zeh, N., Handrick, R., Gorr, I., Schulz, P., Fischer, S., Gamer, M. and Otte, K. (2020) Unraveling what makes a monoclonal antibody difficult-to-express: From intracellular accumulation to incomplete folding and degradation via ERAD. Biotechnol. Bioeng., 117, 5–16.

87. Reinhart, D., Sommeregger, W., Debreczeny, M., Gludovacz, E. and Kunert, R. (2014) In search of expression bottlenecks in recombinant CHO cell lines - A case study. Appl. Microbiol. Biotechnol., 98, 5959–5965.

88. Reisinger, H., Steinfellner, W., Stern, B., Katinger, H. and Kunert, R. (2008) The absence of effect of gene copy number and mRNA level on the amount of mAb secretion from mammalian cells. Appl. Microbiol. Biotechnol., 81, 701–710.

89. Schlatter, S., Stansfield, S.H., Dinnis, D.M., Racher, A.J., Birch, J.R. and James, D.C. (2005) On the optimal ratio of heavy to light chain genes for efficient recombinant antibody production by CHO cells. Biotechnol. Prog., 21, 122–133.

90. Wijesuriya, S.D., Pongo, E., Tomic, M., Zhang, F., Garcia-Rodriquez, C., Conrad, F., Farr-Jones, S., Marks, J.D. and Horwitz, A.H. (2018) Antibody engineering to improve manufacturability. Protein Expr. Purif., 149, 75–83.

91. Pybus, L.P., James, D.C., Dean, G., Slidel, T., Hardman, C., Smith, A., Daramola, O. and Field, R. (2014) Predicting the expression of recombinant monoclonal antibodies in Chinese hamster ovary cells based on sequence features of the CDR3 domain. Biotechnol. Prog., 30, 188–197.

92. Vanhove, M., Usherwood, Y.K. and Hendershot, L.M. (2001) Unassembled Ig heavy chains do not cycle from BiP in vivo but require light chains to trigger their release. Immunity, 15, 105–114.

93. Cartwright, J.F., Arnall, C.L., Patel, Y.D., Barber, N.O.W., Lovelady, C.S., Rosignoli, G., Harris, C.L., Dunn, S., Field, R.P., Dean, G., et al. (2020) A platform for context-specific genetic engineering of recombinant protein production by CHO cells. J. Biotechnol., 312, 11–22.

94. Donaldson, J., Kleinjan, D.J. and Rosser, S. (2022) Synthetic biology approaches for dynamic CHO cell engineering. Curr. Opin. Biotechnol., 78, 102806.

95. Fischer, S. and Otte, K. (2019) CHO Cell Engineering for Improved Process Performance and Product Quality. In Cell Culture Engineering. John Wiley & Sons, Ltd, pp. 207–250.

96. Tadauchi, T., Lam, C., Liu, L., Zhou, Y., Tang, D., Louie, S., Snedecor, B. and Misaghi, S. (2019) Utilizing a regulated targeted integration cell line development approach to systematically investigate what makes an antibody difficult to express. Biotechnol. Prog., 35, e2772.

97. Kaneyoshi, K., Kuroda, K., Uchiyama, K., Onitsuka, M., Yamano-Adachi, N., Koga, Y. and Omasa, T. (2019) Secretion analysis of intracellular “difficult-to-express” immunoglobulin G (IgG) in Chinese hamster ovary (CHO) cells. Cytotechnology, 71, 305–316.

98. Sebastião, M.J., Hoffman, M., Escandell, J., Tousi, F., Zhang, J., Figueroa, B., DeMaria, C. and Gomes-Alves, P. (2023) Identification of Mispairing Omic Signatures in Chinese Hamster Ovary (CHO) Cells Producing a Tri-Specific Antibody. Biomedicines, 11, 2890.

99. Fu, Q., Polanco, A., Lee, Y.S. and Yoon, S. (2023) Critical challenges and advances in recombinant adeno-associated virus (rAAV) biomanufacturing. Biotechnol. Bioeng., 120, 2601–2621.

100. Gutiérrez-Granados, S., Cervera, L., Kamen, A.A. and Gòdia, F. (2018) Advancements in mammalian cell transient gene expression (TGE) technology for accelerated production of biologics. Crit. Rev. Biotechnol., 38, 918–940.

101. Tihanyi, B. and Nyitray, L. (2020) Recent advances in CHO cell line development for recombinant protein production. Drug Discov. Today Technol., 38, 25–34.

102. Ingusci, S., Verlengia, G., Soukupova, M., Zucchini, S. and Simonato, M. (2019) Gene Therapy Tools for Brain Diseases. Front. Pharmacol., 10.

103. Castro, B., Steel, J.C. and Layton, C.J. (2024) AAV-mediated gene therapies for glaucoma and uveitis: are we there yet? Expert Rev. Mol. Med., 26, e9.

104. Hussein, M.K., Papež, M., Dhiman, H., Baumann, M., Galosy, S. and Borth, N. (2022) In silico design of CMV promoter binding oligonucleotides and their impact on inhibition of gene expression in Chinese hamster ovary cells. J. Biotechnol., 359, 185–193.

105. Jalšić, L., Lytvyn, V., Elahi, S.M., Hrapovic, S., Nassoury, N., Chahal, P.S., Gaillet, B. and Gilbert, R. (2023) Inducible HEK293 AAV packaging cell lines expressing Rep proteins. Mol. Ther. Methods Clin. Dev., 30, 259–275.

106. Maltais, J.S., Lord-Dufour, S., Morasse, A., Stuible, M., Loignon, M. and Durocher, Y. (2023) Repressing expression of difficult-to-express recombinant proteins during the selection process increases productivity of CHO stable pools. Biotechnol. Bioeng., 120, 2840–2852.

107. Chen, L., Dickerhoff, J., Sakai, S. and Yang, D. (2022) DNA G-Quadruplex in Human Telomeres and Oncogene Promoters: Structures, Functions, and Small Molecule Targeting. Acc. Chem. Res., 55, 2628–2646.

