## Supplementary Information for "Synthetic G-quadruplex components for predictable, precise two-level control of mammalian recombinant protein expression"

**Supplementary Table S1**. Synthetic G-quadruplex motif sequences. Components utilized in experiments to mechanistically dissect library function and validate performance in bioindustrial applications (i.e. Figs 3-5) are shown in bold. Sequences used as oligonucleotides in CD experiments were flanked by a single thymine at both ends.

| Motif number | Sequence | Component name |
|---|---|---|
| DNA1 | $G_2AG_2CG_2TG_2$ | |
| DNA2 | $G_3AG_3CG_3TG_3$ | |
| DNA3 | $G_4AG_4CG_4TG_4$ | |
| DNA4 | $G_5AG_5CG_5TG_5$ | |
| DNA5 | $G_6AG_6CG_6TG_6$ | |
| DNA6 | $G_2ACG_2CTG_2TAG_2$ | |
| DNA7 | $G_3ACG_3CTG_3TAG_3$ | |
| **DNA8** | **$G_4ACG_4CTG_4TAG_4$** | **DNA.75REU** |
| DNA9 | $G_5ACG_5CTG_5TAG_5$ | |
| **DNA10** | **$G_6ACG_6CTG_6TAG_6$** | **DNA.45REU** |
| **DNA11** | **$G_2ACTG_2CTAG_2TACG_2$** | **DNA.95REU** |
| DNA12 | $G_3ACTG_3CTAG_3TACG_3$ | |
| **DNA13** | **$G_4ACTG_4CTAG_4TACG_4$** | **DNA.70REU** |
| DNA14 | $G_5ACTG_5CTAG_5TACG_5$ | |
| **DNA15** | **$G_6ACTG_6CTAG_6TACG_6$** | **DNA.50REU** |
| DNA16 | $G_2ACTAG_2CTACG_2TACTG_2$ | |
| DNA17 | $G_3ACTAG_3CTACG_3TACTG_3$ | |
| DNA18 | $G_4ACTAG_4CTACG_4TACTG_4$ | |
| DNA19 | $G_5ACTAG_5CTACG_5TACTG_5$ | |
| DNA20 | $G_6ACTAG_6CTACG_6TACTG_6$ | |
| DNA21 | $G_2ACTACG_2CTACTG_2TACTAG_2$ | |
| DNA22 | $G_3ACTACG_3CTACTG_3TACTAG_3$ | |
| DNA23 | $G_4ACTACG_4CTACTG_4TACTAG_4$ | |

| | | |
|---|---|---|
| DNA24 | $G_5ACTACG_5CTACTG_5TACTAG_5$ | |
| **DNA25** | **$G_6ACTACG_6CTACTG_6TACTAG_6$** | **DNA.50REU.2** |
| DNA26 | $G_2ACG_2CTG_2TAG_2ACG_2$ | |
| DNA27 | $G_3ACG_3CTG_3TAG_3ACG_3$ | |
| DNA28 | $G_4ACG_4CTG_4TAG_4ACG_4$ | |
| DNA29 | $G_5ACG_5CTG_5TAG_5ACG_5$ | |
| **DNA30** | **$G_6ACG_6CTG_6TAG_6ACG_6$** | **DNA.30REU** |
| DNA31 | $G_2ACG_2CTG_2TAG_2ACG_2CTG_2$ | |
| DNA32 | $G_3ACG_3CTG_3TAG_3ACG_3CTG_3$ | |
| DNA33 | $G_4ACG_4CTG_4TAG_4ACG_4CTG_4$ | |
| DNA34 | $G_5ACG_5CTG_5TAG_5ACG_5CTG_5$ | |
| **DNA35** | **$G_6ACG_6CTG_6TAG_6ACG_6CTG_6$** | **DNA.20REU** |
| DNA36 | $G_2ACG_2CTG_2TAG_2ACG_2CTG_2TAG_2$ | |
| DNA37 | $G_3ACG_3CTG_3TAG_3ACG_3CTG_3TAG_3$ | |
| DNA38 | $G_4ACG_4CTG_4TAG_4ACG_4CTG_4TAG_4$ | |
| DNA39 | $G_5ACG_5CTG_5TAG_5ACG_5CTG_5TAG_5$ | |
| **DNA40** | **$G_6ACG_6CTG_6TAG_6ACG_6CTG_6TAG_6$** | **DNA.10REU** |
| DNA41 | $G_3AAAG_3AAAG_3AAAG_3$ | |
| DNA42 | $G_3AACG_3AACG_3AACG$ | |
| DNA43 | $G_3AATG_3AATG_3AATG$ | |
| DNA44 | $G_3ACAG_3ACAG_3ACAG$ | |
| DNA45 | $G_3ACCG_3ACCG_3ACCG$ | |
| DNA46 | $G_3ACTG_3ACTG_3ACTG$ | |
| **DNA47** | **$G_3ATAG_3ATAG_3ATAG$** | **DNA.70REU.2** |
| DNA48 | $G_3ATCG_3ATCG_3ATCG$ | |
| DNA49 | $G_3ATTG_3ATTG_3ATTG$ | |
| DNA50 | $G_3CAAG_3CAAG_3CAAG$ | |
| DNA51 | $G_3CACG_3CACG_3CACG$ | |

| | | |
|---|---|---|
| DNA52 | $G_3CATG_3CATG_3CATG$ | |
| DNA53 | $G_3CCAG_3CCAG_3CCAG$ | |
| DNA54 | $G_3CCCG_3CCCG_3CCCG$ | |
| DNA55 | $G_3CCTG_3CCTG_3CCTG$ | |
| DNA56 | $G_3CTAG_3CTAG_3CTAG$ | |
| DNA57 | $G_3CTCG_3CTCG_3CTCG$ | |
| DNA58 | $G_3CTTG_3CTTG_3CTTG$ | |
| DNA59 | $G_3TAAG_3TAAG_3TAAG$ | |
| DNA60 | $G_3TACG_3TACG_3TACG$ | |
| DNA61 | $G_3TATG_3TATG_3TATG$ | |
| DNA62 | $G_3TCAG_3TCAG_3TCAG$ | |
| **DNA63** | $\mathbf{G_3TCCG_3TCCG_3TCCG}$ | **DNA.70REU.3** |
| DNA64 | $G_3TCTG_3TCTG_3TCTG$ | |
| **DNA65** | $\mathbf{G_3TTAG_3TTAG_3TTAG}$ | **DNA.70REU.4** |
| DNA66 | $G_3TTCG_3TTCG_3TTCG$ | |
| DNA67 | $G_3TTTG_3TTTG_3TTTG$ | |
| DNAmut1 | AGTACTAGTCTAAGTTACAGT | |
| DNAmut2 | AAGTTACTAAGTTCTAAAGTTTACAAGTT | |
| **RNA1** | $\mathbf{G_2AG_2CG_2UG_2}$ | **RNA.90REU** |
| **RNA2** | $\mathbf{G_3AG_3CG_3UG_3}$ | **RNA.70REU** |
| RNA3 | $G_4AG_4CG_4UG_4$ | |
| **RNA4** | $\mathbf{G_5AG_5CG_5UG_5}$ | **RNA.25REU** |
| RNA5 | $G_6AG_6CG_6UG_6$ | |
| RNA6 | $G_2ACG_2CUG_2UAG_2$ | |
| **RNA7** | $\mathbf{G_3ACG_3CUG_3UAG_3}$ | **RNA.80REU** |
| RNA8 | $G_4ACG_4CUG_4UAG_4$ | |
| **RNA9** | $\mathbf{G_5ACG_5CUG_5UAG_5}$ | **RNA.50REU** |
| RNA10 | $G_6ACG_6CUG_6UAG_6$ | |

| | | |
|---|---|---|
| **RNA11** | **$G_2ACUG_2CUAG_2UACG_2$** | **RNA.90REU** |
| RNA12 | $G_3ACUG_3CUAG_3UACG_3$ | |
| **RNA13** | **$G_4ACUG_4CUAG_4UACG_4$** | **RNA.65REU** |
| RNA14 | $G_5ACUG_5CUAG_5UACG_5$ | |
| **RNA15** | **$G_6ACUG_6CUAG_6UACG_6$** | **RNA.40REU** |
| RNA16 | $G_2ACUAG_2CUACG_2UACUG_2$ | |
| RNA17 | $G_3ACUAG_3CUACG_3UACUG_3$ | |
| RNA18 | $G_4ACUAG_4CUACG_4UACUG_4$ | |
| RNA19 | $G_5ACUAG_5CUACG_5UACUG_5$ | |
| **RNA20** | **$G_6ACUAG_6CUACG_6UACUG_6$** | **RNA.35REU** |
| RNA21 | $G_2ACUACG_2CUACUG_2UACUAG_2$ | |
| RNA22 | $G_3ACUACG_3CUACUG_3UACUAG_3$ | |
| RNA23 | $G_4ACUACG_4CUACUG_4UACUAG_4$ | |
| RNA24 | $G_5ACUACG_5CUACUG_5UACUAG_5$ | |
| RNA25 | $G_6ACUACG_6CUACUG_6UACUAG_6$ | |
| RNA26 | $G_2ACG_2CUG_2UAG_2ACG_2$ | |
| RNA27 | $G_3ACG_3CUG_3UAG_3ACG_3$ | |
| RNA28 | $G_4ACG_4CUG_4UAG_4ACG_4$ | |
| RNA29 | $G_5ACG_5CUG_5UAG_5ACG_5$ | |
| **RNA30** | **$G_6ACG_6CUG_6UAG_6ACG_6$** | **RNA.15REU** |
| RNA31 | $G_2ACG_2CUG_2UAG_2ACG_2CUG_2$ | |
| RNA32 | $G_3ACG_3CUG_3UAG_3ACG_3CUG_3$ | |
| RNA33 | $G_4ACG_4CUG_4UAG_4ACG_4CUG_4$ | |
| RNA34 | $G_5ACG_5CUG_5UAG_5ACG_5CUG_5$ | |
| RNA35 | $G_6ACG_6CUG_6UAG_6ACG_6CUG_6$ | |
| RNA36 | $G_2ACG_2CUG_2UAG_2ACG_2CUG_2UAG_2$ | |
| **RNA37** | **$G_3ACG_3CUG_3UAG_3ACG_3CUG_3UAG_3$** | **RNA.50REU** |
| RNA38 | $G_4ACG_4CUG_4UAG_4ACG_4CUG_4UAG_4$ | |

| | | |
|---|---|---|
| RNA39 | $G_5ACG_5CUG_5UAG_5ACG_5CUG_5UAG_5$ | |
| **RNA40** | **$G_6ACG_6CUG_6UAG_6ACG_6CUG_6UAG_6$** | **RNA.5REU** |
| RNA41 | $G_3AAAG_3AAAG_3AAAG_3$ | |
| RNA42 | $G_3AACG_3AACG_3AACG$ | |
| RNA43 | $G_3AAUG_3AAUG_3AAUG$ | |
| RNA44 | $G_3ACAG_3ACAG_3ACAG$ | |
| RNA45 | $G_3ACCG_3ACCG_3ACCG$ | |
| RNA46 | $G_3ACUG_3ACUG_3ACUG$ | |
| RNA47 | $G_3AUAG_3AUAG_3AUAG$ | |
| RNA48 | $G_3AUCG_3AUCG_3AUCG$ | |
| RNA49 | $G_3AUUG_3AUUG_3AUUG$ | |
| RNA50 | $G_3CAAG_3CAAG_3CAAG$ | |
| RNA51 | $G_3CACG_3CACG_3CACG$ | |
| RNA52 | $G_3CAUG_3CAUG_3CAUG$ | |
| RNA53 | $G_3CCAG_3CCAG_3CCAG$ | |
| RNA54 | $G_3CCCG_3CCCG_3CCCG$ | |
| RNA55 | $G_3CCUG_3CCUG_3CCUG$ | |
| RNA56 | $G_3CUAG_3CUAG_3CUAG$ | |
| RNA57 | $G_3CUCG_3CUCG_3CUCG$ | |
| RNA58 | $G_3CUUG_3CUUG_3CUUG$ | |
| RNA59 | $G_3UAAG_3UAAG_3UAAG$ | |
| RNA60 | $G_3UACG_3UACG_3UACG$ | |
| RNA61 | $G_3UAUG_3UAUG_3UAUG$ | |
| RNA62 | $G_3UCAG_3UCAG_3UCAG$ | |
| RNA63 | $G_3UCCG_3UCCG_3UCCG$ | |
| RNA64 | $G_3UCUG_3UCUG_3UCUG$ | |
| RNA65 | $G_3UUAG_3UUAG_3UUAG$ | |
| RNA66 | $G_3UUCG_3UUCG_3UUCG$ | |

| | |
|---|---|
| RNA67 | $G_3UUUG_3UUUG_3UUUG$ |
| RNAmut1 | AGUACUAGUCUAAGUUACAGU |
| RNAmut2 | AAGUUACUAAGUUCUAAAGUUUACAAGUU |

**Supplementary Table S2.** Sequences of endogenous G4-quadruplex motifs, and their mutated versions (G-to-A mutations underlined). Propensity of sequences to form G4 structures was predicted using QGRS mapper (49).

| Motif | Sequence | G-score |
|---|---|---|
| c-myc wild type | TGGGGAGGGTGGGGAGGGTGGGGAAGG | 41 |
| c-myc mut1 | TGAGGAGAGTGAGGAGAGTGAGGAAGG | 16 |
| c-myc mut2 | TGAAGAGAATGAAGAGAATGAAGAAGG | 0 |
| Nras wild type | GGGAGGGGCGGGUCUGGG | 40 |
| Nras mut1 | AAAAGGGGCGGGUCUGGG | 18 |
| Nras mut2 | AAAAGAGGCGAGUCUGAG | 0 |
